# High-throughput sequencing of murine immunoglobulin heavy chain transcripts using single side unique molecular identifiers on an Ion Torrent PGM

**DOI:** 10.1101/219568

**Authors:** Jean-Philippe Bürckert, William J. Faison, Axel R. S. X. Dubois, Regina Sinner, Oliver Hunewald, Anke Wienecke-Baldacchino, Anne Brieger, Claude P. Muller

**Author notes:** Corresponding author Mailing address: Luxembourg Institute of Health, House of Biohealth, 29, rue Henri Koch, 4354 Eschsur-Alzette, Luxembourg J.-P. Bürckert - C. P. Muller –.

## Abstract

With the advent of high-throughput sequencing (HTS), profiling immunoglobulin (IG) repertoires has become an essential part of immunological research. Advances in sequencing technology enable the IonTorrent Personal Genome Machine (PGM) to cover the full-length of IG mRNA transcripts. Nucleotide insertions and deletions (indels) are the dominant errors of the PGM sequencing platform and can critically influence IG repertoire assessments. Here, we present a PGM-tailored IG repertoire sequencing approach combining error correction through unique molecular identifier (UID) barcoding and indel detection through ImMunoGeneTics (IMGT), the most commonly used sequence alignment database for IG sequences. Using artificially falsified sequences for benchmarking, we found that IMGT efficiently detects 98% of the introduced indels through gene-segment frameshifts. Undetected indels are either located at the ends of the sequences or produce masked frameshifts with an insertion and deletion in close proximity. IMGT’s indel correction algorithm resolves up to 87% of the tested insertions, but no deletions. The complementary determining regions 3 (CDR3s) are returned 100% correct for up to 3 insertions or 3 deletions through conservative culling. We further show, that our PGM-tailored unique molecular identifiers results in highly accurate HTS datasets if combined with the presented data processing. In this regard, considering sequences with at least two copies from datasets with UID families of minimum 3 reads result in correct sequences with over 99% confidence. The protocol and sample processing strategies described in this study will help to establish benchtop-scale sequencing of IG heavy chain transcripts in the field of IG repertoire research.

## Introduction

The diversity of the immunoglobulin (IG) repertoire is the key feature of the adaptive immune system, enabling it to theoretically combat every possible antigen encountered during an individual’s lifetime [1]. With the development of high-throughput sequencing (HTS) it became possible to analyze the IG repertoire at high depth [2–6]. Studies, almost a decade ago, established Roche’s 454 sequencer as the first tool of choice for exhaustive characterization of IG repertoires due to its superior read-length [7]. More recently, Illumina’s MiSeq and HiSeq sequencers as well as the Ion Torrent Personal Genome Machine (PGM, Thermo Fisher Scientific) provided an improved sequencing technologies which can reach across the full V(D)J nucleotide sequence span [8]. The different technologies of the sequencers result each in their specific error-rates and -types [7,9–15]. Illumina’s optical sequencing produces mostly nucleotide (nt) transversions and transitions, which can be corrected by building consensus sequences [16]. The 454’s pyrosequencing chemistry and the PGMs semiconductor technique mainly introduce homopolymer repeats resulting in insertions and deletions of bases, which can be corrected by gene segment-wise reference alignment [17].

Most sequencing approaches use IG isotype specific constant (C) region primers to translate IG heavy-chain (IGH) (m)RNA into cDNA, which are subsequently amplified using a set of V-region specific primers in a multiplex PCR approach. However, this can result in skewed repertoire read-outs due to biased PCR efficacy [8,14,18]. In addition, sequencing errors can falsify somatic hypermutation profiles, VDJ germline gene assignment and clonal grouping [8,19]. Unique identifiers (UID) which tag individual RNA molecules at cDNA transcription level have been used to obtain an unbiased view on the IG repertoire [20–23]. This method also allows thorough error-correction by building consensus sequences, albeit at the cost of sequencing depth. In all cases, complex bioinformatic approaches are necessary to perform raw-read processing [24]. Subsequent alignments to germline genes to assign VDJ family genes are in general conducted using the ImMunoGeneTics (IMGT) database, which applies an error correction algorithm for insertions and deletions in the process [25,26].

After the initial proof-of-concept studies, the use of animal models to study the IG repertoire dynamics has been largely ignored [4,6]. One major factor being the lack of a suitable IGH V-region primer set comparable to BIOMED-2, developed for the human IG repertoire [27]. Yet, animal models offer advantages over human studies, as they are not limited to peripheral blood and have a lower B cell diversity [28–31]. As IMGT provides repertoires for various species, we chose to develop a method to profile the IG repertoire of Balb/C mice, one of the most commonly used animal models.

In the present study, the performance of the PGM sequencing platform together with the IMGT database for the assessment of murine IGH repertoires is evaluated. In this context, several novel aspects are examined: first, the IMGT database’s indel detection and correction algorithm is benchmarked with a set of artificially falsified sequences. Second, a 16-nucleotide single side UID (ssUID) barcoding technique tailored to the PGM sequencing chemistry is introduced together with a swift 1-day library preparation protocol. Third, the PGM’s error-rate for sequencing murine IG transcripts with our barcoding strategy and customized data processing is determined.

## Results

### Reference sequences

A set of 7 monoclonal mouse hybridoma cell lines was used to investigate the distribution and influence of insertions and deletions (indels) produced by the IonTorrent PGM sequencing technology on murine IGH repertoire sequencing (Figure 1). Reference sequences were obtained from Sanger sequenced cDNA transcripts of monoclonal hybridoma RNA subsequently annotated and translated into amino acids by IMGT V-QUEST.

**Figure 1:**
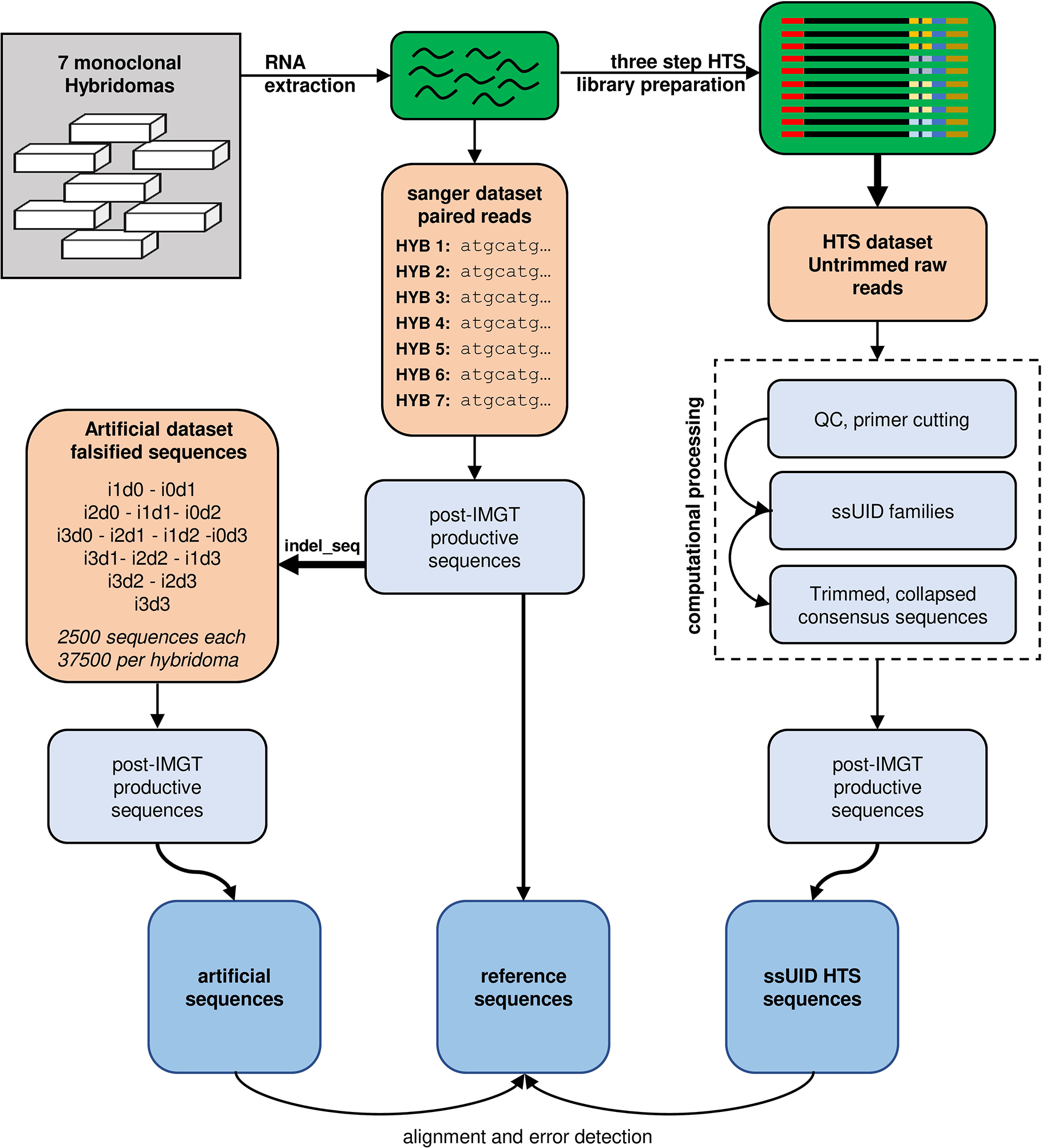
Study design. RNA was extracted from 7 monoclonal hybridoma cell lines and reverse transcribed into cDNA. cDNA sequences were determined by Sanger sequencing and submitted to IMGT to determine reference sequences. Reference sequences were artificially falsified using the indel_seq program, introducing up to 3 insertions and 3 deletions. 2500 artificial sequences were generated for each permutation and hybridoma and processed by IMGT. Post-IMGT sequences were aligned to the references to determine error detection and correction. RNA was also used to generate high-throughput sequencing (HTS) libraries in a three-step library preparation protocol. Single side unique identifiers (ssUID) were introduced during reverse transcription to tag each RNA molecule individually (see also **suppl. Fig. S3**). Libraries were sequenced on an Ion Torrent PGM sequencer with all quality trimming options disabled in the Torrent Suite software. Untrimmed raw sequences were processed with a custom-made bioinformatics pipeline generating consensus sequences per UID family. Collapsed consensus sequences were submitted to IMGT and post-IMGT sequences aligned to the reference sequences to determine error detection and correction.

### Distribution of artificial insertions and deletions

To investigate the influence of indels on IMGT processing of an IGH sequence, we generated a benchmark dataset from the reference sequences that contained artificially introduced indels at random positions (**suppl. table S1**). To cover each position within a 300 nt sequence with minimum 90% certainty, at least 2398 erroneous variants are required [32]. Therefore, we generated 2500 artificial, randomly flawed sequences for each permutation of 0-3 insertions and/or deletions (indels, annotated as i1d0, i0d1, i1d1…i3d3), resulting in a total of 37500 artificial sequences per original hybridoma sequence with indels ranging from 1 to 6 events. Indels were homogenously present as determined by graphical reference alignment (Fig. 2A). Uncovered positions resulted from indels within homopolymer stretches which were always assigned to the beginning of such a nucleotide repeat region (Fig. 2B).

**Figure 2:**
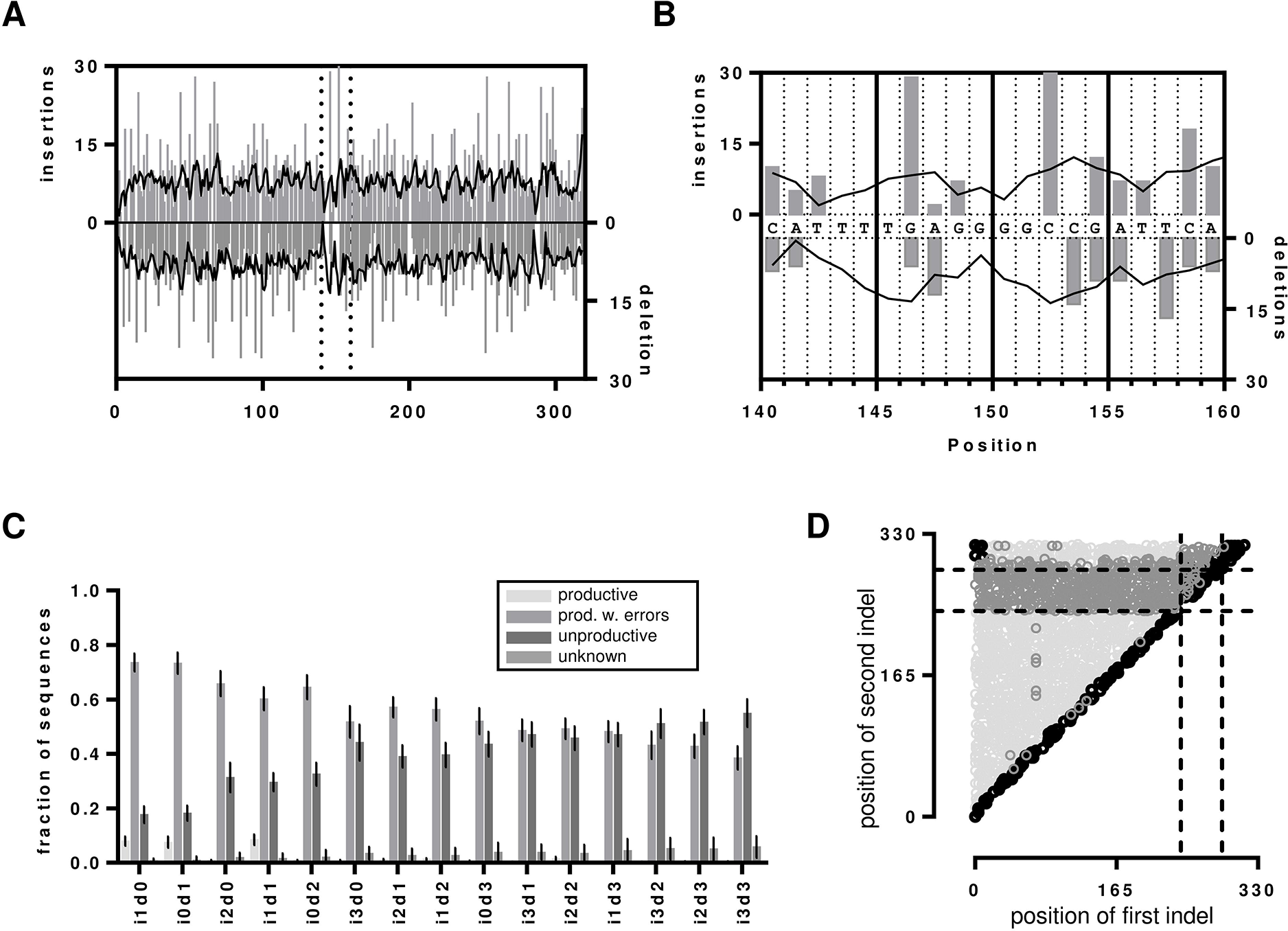
Indels in the artificial dataset. (A) Insertion and deletion events displayed as determined by graphical alignments of the reference sequence to the i1d0 and i0d1 dataset of hybridoma 1. Grey bars represent the actual detected indel and the black line presents the moving average over 4 neighbors. The dotted lines vertical present the segment that is magnified in (B) to visualize the problem of determining the position of indels in homopolymer repeats. (C) Indel detection rates by IMGT processing shown as bar chart with error bars indicating the SD over all 7 datasets (D) Visualization of indel proximity. The distances between the first and second indel before correction in the i1d1 dataset of hybridoma 1 are shown as scatterplot. Dotted lines indicate the position of the IMGT junction. Productive sequences with detected indels are shown in light grey, unproductive sequences are shown in dark grey. Sequences without detected errors are shown in black. The remaining i1d1 indel proximity graphs are shown in the **supplementar Figure S1.**

### IMGT VDJ nt error detection

As each sequence of the benchmark system contained indel errors, all sequences marked by IMGT as productive were falsely categorized as error free. In general, IMGT correctly recognized 97.9% (± 2.9%) of the introduced indels over all datasets and categorized the sequences then either as productive with detected indels, unproductive or unknown (Fig. 2C). Interestingly, only the sets with one insertion and/or deletion (i1d0, i0d1 and i1d1) exhibited elevated numbers of unrecognized indels. For these IMGT falsely returned 8% (±1.8%) of the sequences as productive, whereas for all other datasets it was only 0.7% (± 0.4%). Such undetected indels were found at the beginning and the end of the sequence or across the whole sequence for i1d1 datasets due to indels in close proximity to each other masking the frame-shifts (Fig. 2D, Fig. 3, **suppl. Fig. S1 and S2**). The number of unproductive sequences increased with the number of indel events, regardless of their composition. Accordingly, the number of productive sequences with detected indels decreased. Less than 50% of sequences with more than 3 indels, were retained. Indels were homogenously distributed in the uncorrected productive sequences with detected errors until about 4/5^th^ of the sequence lengths while the opposite is true for the uncorrected unproductive sequences (Fig. 2D, Fig. 3 **and suppl. Fig. S2**). This section of the sequence coincides with the IMGT IGH junction which encodes for the CDR3 [33]. Accordingly, upon detecting an indel in the IGH junction, IMGT categorized the sequence as unproductive and no corrective attempts were made.

**Figure 3:**
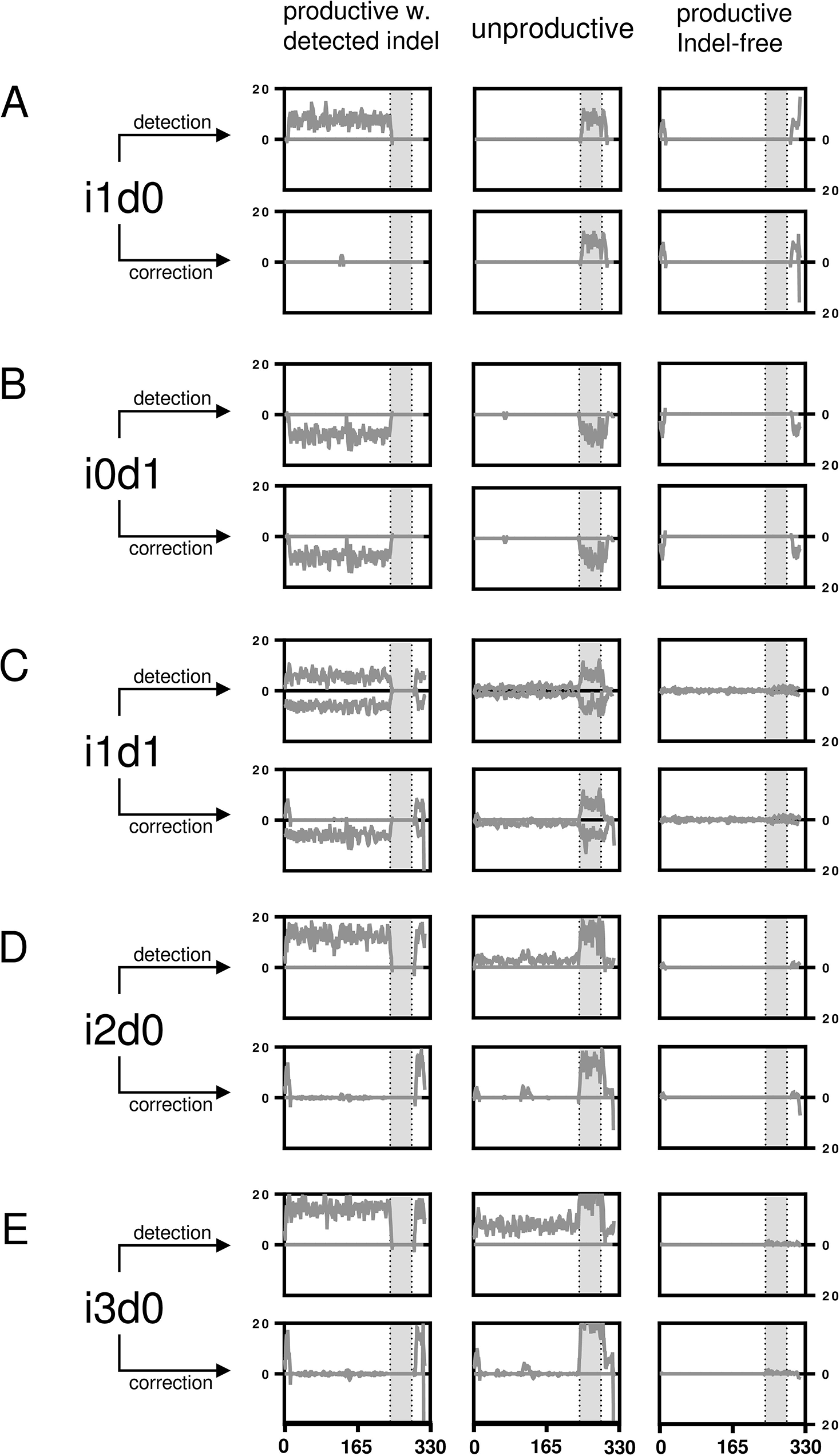
Artificial indel set alignments. Indel positions are shown before and after IMGT error correction for artificially falsified Hybridoma 1 sequences separated by productivity. (A) The indels for the i1d0 dataset are shown per nucleotide position as line plot (smoothened over 4 neighbors). The grey area marks the IGH VDJ junction. (B-E) like (A) but with different permutations. The remaining permutations are displayed in the **supplementary Figure S2.**

### Nucleotide error correction

Upon detection of an indel, IMGT tries to correct it by alignment to its closest germline. The efficacy of this process was investigated by aligning the sequences with detected indels to determine the number of correctly resolved sequences (Fig. 3, Fig. 4 **and suppl. Fig. S2**). A thorough error reduction was observed for up to three insertion errors in datasets without deletions, returning 87% ± 3.2% (i1d0), 72% ± 5.5% (i2d0) and 56% ± 7.0% (i3d0) of productive sequences as correct (Fig. 4). Within these sequences indels that were not corrected by the IMGT were mainly found at the beginning and end of the sequence (Fig. 3A, D, E). In the case of deletions, the IMGT correction introduced a gap for the missing nucleotide as the original nucleotide was unknown. Consequently, the number of correct sequences found in datasets with mixed insertions and deletions is very low (i1d1: 1% ± 0.3%, i2d1: 2% ± 0.3%, i3d1: 2% ± 0.6%, i2d2 and i3d2 <1%). Nevertheless, in these datasets, the insertions within the sequences were always reduced (Fig 3C **and suppl. Fig. S2**). No correct sequence could be identified in deletion-only datasets (Fig. 4).

**Figure 4:**
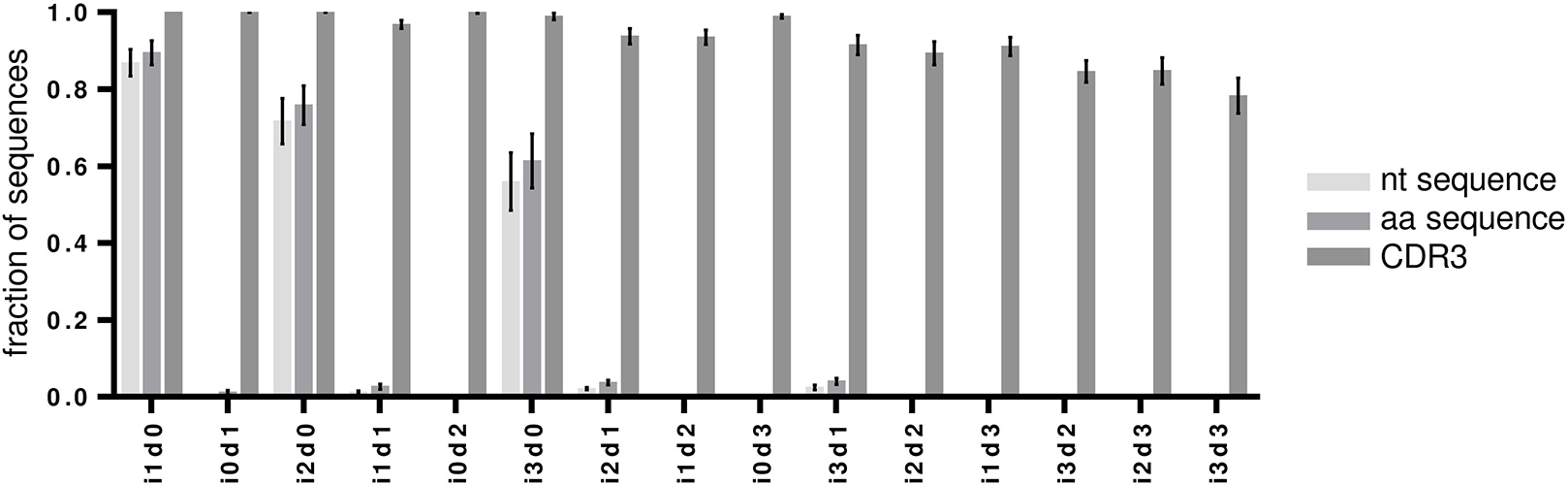
Correction of artificially introduced indels by IMGT. The fraction of correct sequences for each artificial benchmark permutation of indels are shown as bar charts of nucleotide (nt), amino acid (aa) and CDR3 amino acid sequences. Error bars indicate SD over all 7 datasets.

### Amino acid error correction

Theoretically, translated amino acids are less influenced by sequencing errors because of the redundancy of the genetic code. Thus, most amino acid translations were returned correctly in the case of insertion-only datasets and with slightly higher numbers compared to the nucleotide datasets (mean correct amino acid sequences for i1d0: 89% ± 2.9%, i2d0: 76% ± 4.7%, i3d0: 61% ± 6.5%, Fig. 4). Higher numbers of correct translations were observed in mixed indel datasets than for the corresponding nucleotide datasets (i1d1: 3% ± 0.7%, i2d1: 4% ± 0.6%, i3d1: 4% ± 0.8%, i2d2 and i3d2 <1%, Fig. 4). Interestingly, some amino acid translations were found to be correct for the i0d1 datasets (1% ± 0.5%, **Fig. 4**). Deletion-affected datasets were usually returned with the wrong amino acid sequence by the IMGT algorithm. During IMGT processing, nucleotide deletions rendered the whole codon triplet elusive and were translated as gaps in the amino acid sequence.

Remarkably, the CDR3 proved to be protected chiefly from insertions and deletions through a more conservative correction approach of the IMGT algorithm for this part of the sequence. As mentioned above, detected indels within the IGH junction, and thus the CDR3, corrupted the entire sequence as unproductive (Fig. 3 **and suppl. Figure S2**). Culling attempts by IMGT turned out to be largely successful (100% correct CDR3s for up to 3 insertions or 3 deletions). Even for the i3d3 indel permutation, IMGT returned 78% ± 4.3% correct CDR3s (Fig 4), by removing all those sequences where indels were detected in the CDR3 encoding nucleotides. Datasets with simultaneous insertions and deletions showed in general lower numbers of correct CDR3 sequences (range 78-97%). This resulted from sequences where indels were introduced in close proximity of each other, producing no detectable frameshift within the IGH junction (Fig 2D). While invisible for the IMGT algorithm, they were observed as variants of the correct CDR3 amino acid sequence.

Taken together the above data show, that IMGT processing exhibits adequate detection of indels through frame-shifts in mouse IGH nt sequences. Consequently, frame-shift masking error compositions cannot be detected and result in amino acid changes in the translations. IMGTs indel correction proved to be reliable for single insertions. However, the impossibility to correct for deletions and larger indel permutations makes consideration of sequences categorized as “productive with detected indels” unfavorable.

### HTS of hybridoma ssUID libraries

Next, the IMGT database and a PGM-tailored data processing pipeline developed by our group were tested using real HTS datasets derived from 7 monoclonal hybridomas (**Figure 1**). The HTS libraries were prepared using an IonTorrent PGM tailored single-side UID approach (**suppl. Fig. S3**) allowing for error correction through building consensus sequences from all reads within a UID family [34,35]. The ssUID barcodes together with the C-region primer and appropriate ‘GATC’ spacer were correctly identified at the sequencing start site of 99.12% ± 0.56% of the usable reads containing a sample specific MID (Table 1). Between 146,010 and 739,854 reads were obtained per sample, with varying ssUID family size distributions (**Fig. 5A**). After raw data processing, 1,431 to 47,169 consensus sequences were retained per hybridoma (Table 1) and uploaded to IMGT HighV-QUEST.

**Table 1.**
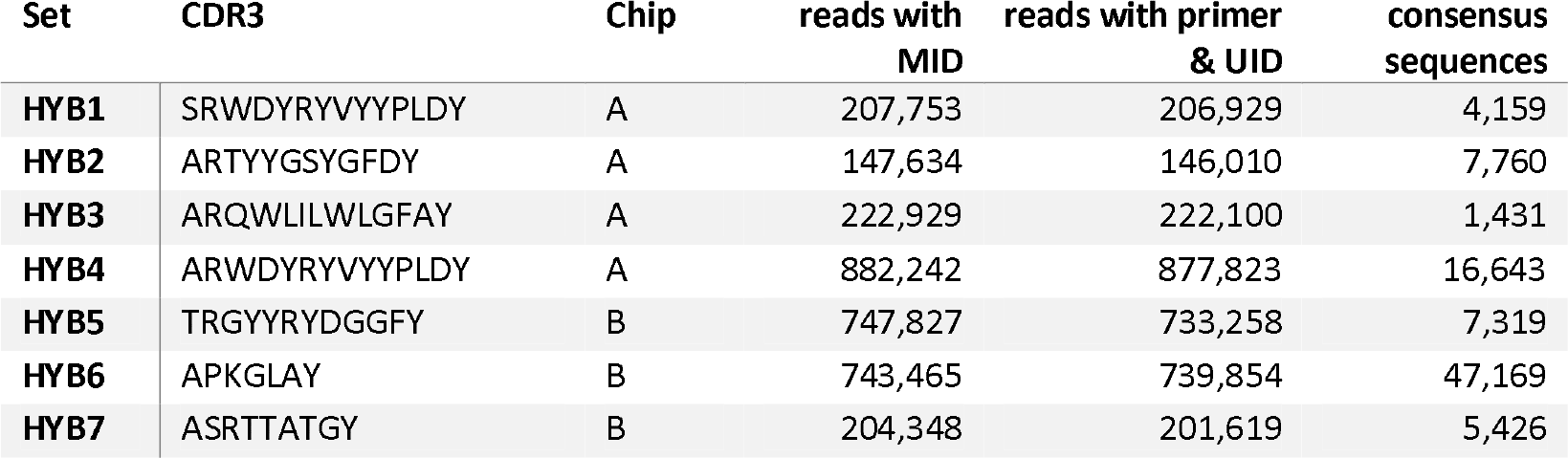
HTS datasets pre-IMGT.

**Figure 5:**
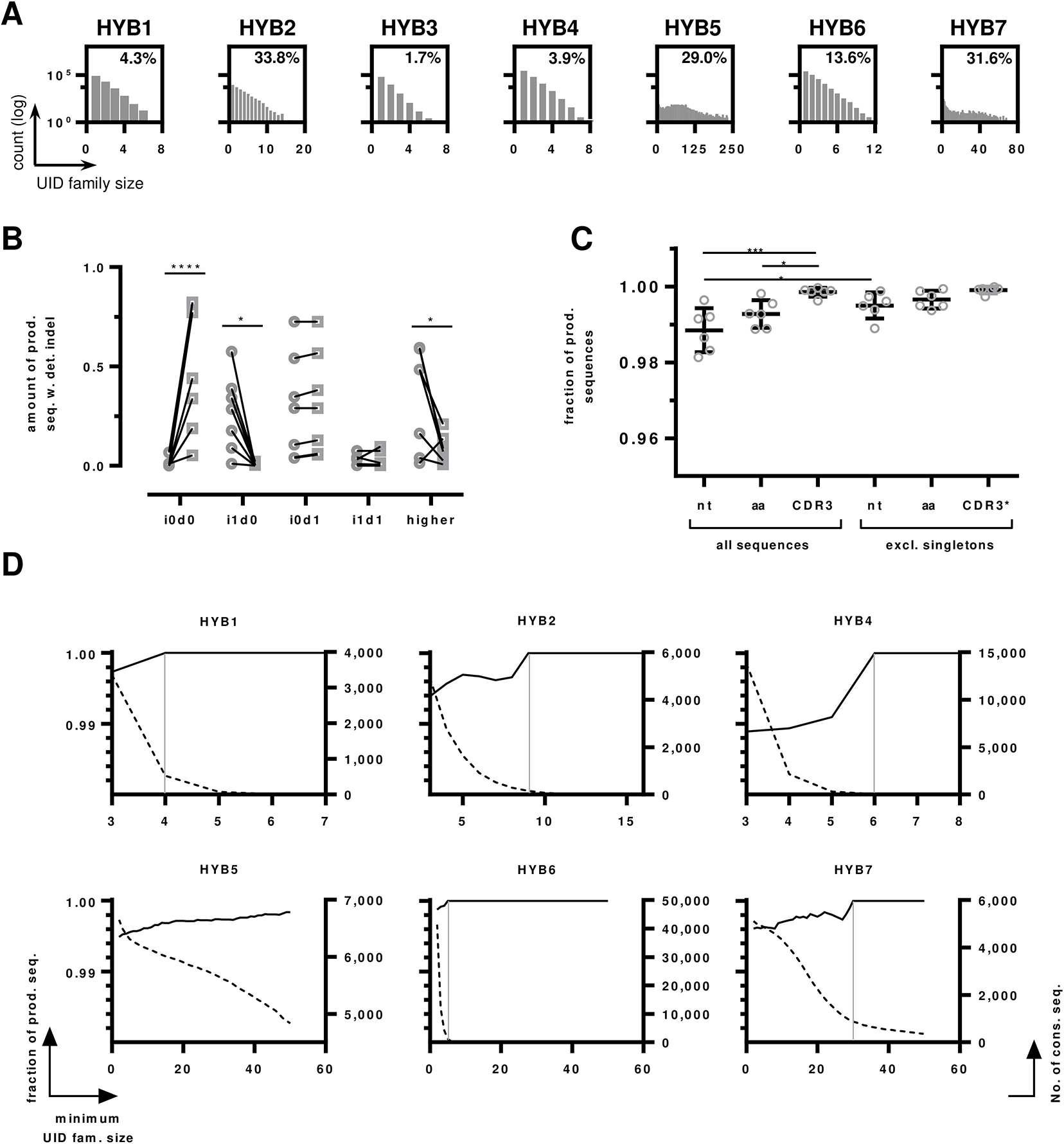
HTS data on monoclonal hybridomas. (A) UID family size distributions per sample. The number of UID families (log transformed) is plotted by the number of reads assigned to a ssUID per hybridoma. The amount of UID families containing a minimum of 3 reads are indicated as percentage value. (B) Indel distributions on productive sequences with detected errors. The amount of indel-free (i0d0), single insertions (i1d0), single deletions (i0d1), one single insertion and deletion (i1d1) and higher permutations are shown as fraction of productive reads with detected indels before (circles) and after (squares) IMGT error correction. Statistical differences are indicated with **** p < 0.0001, * p < 0.05, multiple two tailed t-test with Holm-Sidak’s method to account for multiple testing. (C) The number of error-free sequences in the productive dataset without detected indels are shown as scatterplot with mean and ± SD. Data are shown for all nucleotide sequences (nt), amino acid sequences (aa) and CDR3s for all sequences and data without singleton sequences. CDR3 singleton exclusion was performed on the basis of full-length amino acid sequences. P values are indicated *** p < 0.001, * < 0.05, One-way ANOVA with Sidak’s post-hoc test. All other differences were not statistically significant. (D) Influence of UID family size on the number of correct sequences. The number of correct sequences are shown as black line per minimum UID family size (left y-axis). The number of consensus sequences are shown as dotted line per minimum family size (right y-axis). The UID family size at which all sequences are correct is indicated by a grey vertical line for Hybridoma 1,2,4,6 and 7, the dataset of Hybridoma 5 does not reach 100% correct sequences.

### IMGT processing of HTS datasets

The majority of the post-IMGT sequences were categorized as productive (75.8% ± 22.6%) and 10.9% (± 9.6%) were categorized as productive with detected indels (Table 2). The remaining sequences were either categorized as unproductive or unknown/else. To investigate the undetected or uncorrected errors within the two productive categories, sequences were aligned to their corresponding references. For Hybridoma 3, which had the poorest UID distribution (**Figure 5A**), only 26.8% of the sequences were classified as productive and 68.8% unproductive (Table 2). This hybridoma was therefore excluded from further analysis.

**Table 2.**
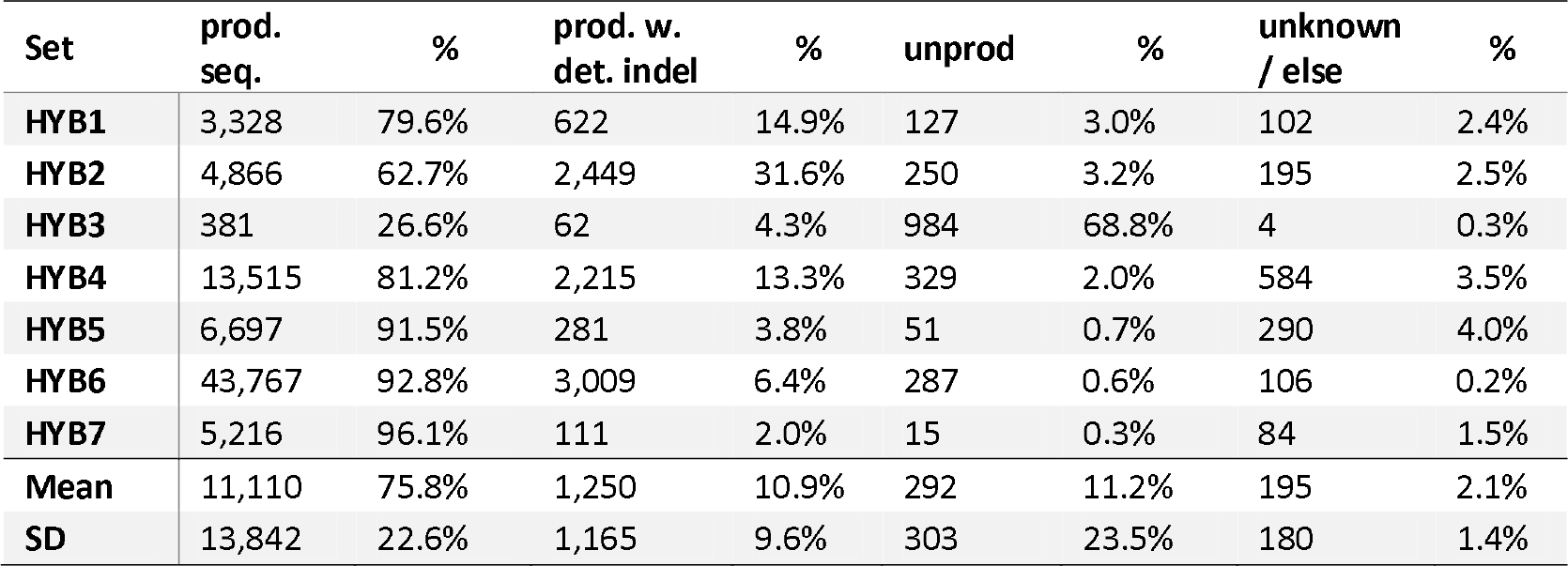
HTS datasets post-IMGT.

In the group of productive sequences with detected errors, IMGT’s indel correction algorithm improved the number of correct sequences by 54.1% to on average 55.3% (± 32.0%, **Fig. 5B**). As expected, IMGT corrected most sequences that contained single insertions efficiently, reducing these errors from average 25.2 (± 24.3%) to 0.48% (± 0.72%, p-value = 0.0027, two-tailed t-test in Graphpad Prism, using Holm-Sidak’s method [36] to account for multiple testing with alpha = 5%, **Figure 5B**). Single deletions were found at somewhat higher rates than single insertions (29.9% ± 24.3%) of the sequences. They increased slightly after IMGT error correction (31.6% ± 24.1%), as insertions of higher indel permutations were corrected, leaving only deletions in the sequences. Accordingly, these higher permutations were found in 33.8% (± 23.8%) of the sequences before error-correction and reduced to 8.8% (± 6.3%) afterwards. While the detection of indel errors in the sequences by IMGT was efficient, the remaining errors after correction still affected 44.7% ± 32.2% of the sequences. As described for the benchmarking sequences above, makes further consideration of sequences marked as “productive with detected indels” inadvisable.

Sequences marked as productive without detected indels are not modified by IMGT but can nonetheless contain indel and nucleotide substitution errors. IMGT does not detect ambiguous nucleotides as errors but marks them as silent mutations. On average 2.2% (± 1.6%) of the consensus sequences in the productive dataset without detected indels contained ambiguous nucleotides (Table 3), which were discarded from the datasets. Most of the remaining sequences were indeed error-free (98.8% ± 0.5%, **Fig. 5C**). The other 1.2% contained on average 0.2% (± 0.1%) i1d1 indels in close proximity to each other, masking frameshifts. Some sequences showed single insertions (0.1% ±0.2%) and deletions (0.15% ± 0.13%), either at the beginning or the end, without detectable frameshift. The remaining false sequences contained nucleotide substitutions, with the majority being transversions (0.5% ± 0.3%) and very few transitions (< 0.1%). As described by Shugay and coworkers, such substitutions originate from dominating polymerase errors occurring early during the amplification [34]. As polymerase errors are occurring at relatively random positions, it is stochastically unlikely, that the same errors are found repeatedly within a dataset and can thus be accounted for by considering only consensus sequences that appear more than once in the final dataset [34,35]. Following this approach, the data was reassessed, excluding singleton consensus sequences. This reduced the number of total sequences in the datasets by 0.8% (± 0.4%). The number of transversions was reduced significantly by 0.3% to 0.16% (± 0.19%, p-value = 0.008, two--tailed t-test in Graphpad Prism, using Holm-Sidak’s method to account for multiple testing with alpha = 5%, data not shown). Consequently, the number of error-free sequences improved significantly by 0.7% to 99.5% (± 0.3%, p-value < 0.0001, two-tailed t-test, using Holm-Sidak’s method to account for multiple testing with alpha = 5%).

**Table 3.**
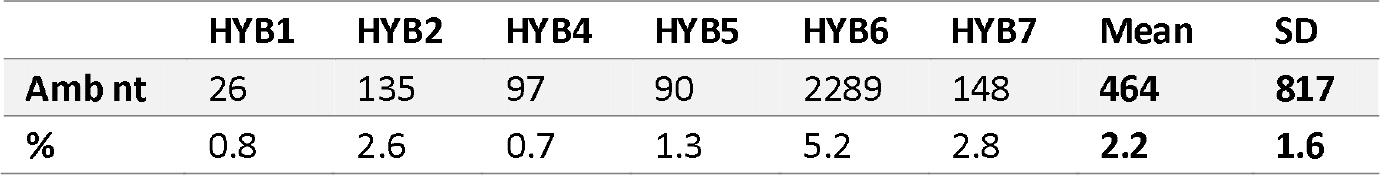
Ambiguous nt in HTS datasets.

The number of reads per UID, referred to as UID family size, is crucial to obtain reliable consensus sequences [35]. Increasing the minimum number of required reads per UID family improved the amount of correct sequences, reaching 100% for all hybridomas, except Hybridoma 5, albeit with different UID family sizes (**Figure 5D**). However, with increasing minimum UID family sizes, the number of sequences decreased exponentially. Consequently, at the point of reaching 100% correct sequences, on average only 7.9% (± 7.1%, excl. Hybridoma 5) of the sequences remained (**Figure 5D**). According to our data, keeping a minimum UID family size of 3 provided adequate accuracy and throughput when using an IonTorrent PGM.

As expected, the number of correct amino acid sequences was higher (99.3% ± 0.3%) than the amount of correct nucleotide sequences (**Figure 5C**). An average of 0.6% (± 0.4%) of the sequences was subject to amino acid changes. Excluding singleton amino acid sequences increased the number of correct amino acid sequences to 99.7% (± 0.2%), but this increase was not statistically significant. CDR3 amino acid sequences were returned almost entirely correct (99.85% ± 0.11%, Figure 31C), increasing to 99.91% (± 0.08%) when singleton full-length amino acid sequences were excluded.

## Discussion

Investigation of IG repertoires by HTS is challenging both with respect to the library preparation as well as sequencing error assessment and data processing. Using artificially falsified sequences, we show here that the IMGT indel detection algorithm is efficient while the IMGT indel correction algorithm only corrects single insertions sufficiently. We confirm the utility of the IonTorrent PGM to assess murine IGH repertoires with high confidence, using a dedicated library preparation protocol with a PGM-tailored 16 nt single side unique identifier (ssUID) barcoding technique. Our data show, that appropriate data processing reduced the error rate of PGM-sequenced IGH repertoires to less than 0.5% false nucleotide and amino acid sequences, and to less than 0.01% false CDR3 sequences per dataset.

Sequencing of IGH repertoires requires a thorough assessment and correction of platform inherent sequencing errors [7,9,12–15]. Using the IMGT database for reference alignment, the indel errors of the utilized Ion Torrent PGM sequencing platform can theoretically be detected through the resulting codon frame-shifts [17]. The VDJ structure of the IGH sequence facilitates indel detection by frame-shift, since gene segments can be aligned separately. In our study, the IMGT algorithm successfully detects 97.9% of all indels, regardless of their composition, only single insertions or deletions at the beginning or the end of the sequences (7.9% and 7.5%, respectively), or i1d1 compositions in close proximity to each other could not be identified (8.5%). IMGT tries to correct detected insertions subsequently by removing the false nucleotide(s) according to the predicted germline sequence. In the artificially falsified datasets of our study insertion-only errors were corrected by the IMGT algorithm with 87% (i1d0), 72% (i2d0) and 56% (i3d0) efficiency. Deletions, on the other hand, are more difficult to recover since the missing nucleotide cannot necessarily be inferred from the germline sequence with sufficient confidence. Consequently, artificially introduced deletions were not corrected by IMGT. Also, for sequences with mixed insertions and deletions only the nucleotide insertions were corrected by IMGT leaving the sequence erroneous. Taken together, these data indicate that detection of indels by IMGT is highly efficient and sequences categorized as “productive” without detected errors are almost entirely indel-free. The low efficiency of the indel correction algorithm makes it inadvisable to take productive sequences with detected indels into account for any downstream analysis. These correspond to about 10% of the final HTS consensus sequences in our study.

HTS library preparation using multiple primers during template amplification can significantly bias the repertoire composition [14,19]. This bias is essentially removed by UID barcoding but the approach reduces sequencing depth at the same time [35,37–39]. In our study, the raw sequencing depth does not influence the relative number of correct sequences while the average UID family size proved to be crucial. For instance, Hybridoma 3, although having only the 3^rd^ lowest amount of raw-reads, lacked eligible UID family sizes (> 2 sequences per UID). For this Hybridoma 3, less than 0.5% of the consensus sequences were built from UID families with more than 2 members, resulting in the poorest error correction rate during sample processing. Consequently, IMGT returned only 26.6% of the consensus sequences as productive. We therefore conclude from our data, that for applying a UID family-wise consensus building approach, samples with less than 0.5% eligible consensus reads after pre-IMGT processing do not have enough coverage to achieve sufficient confidence and depth for the post-IMGT sequences and should be discarded from further analysis.

For grouping reads by UID families, it is essential to identify the UID tags correctly [35,39]. The PGM sequencing chemistry is unidirectional, starting with the sequencing adapter A. Comparable protocols for the Illumina sequencing platforms usually consist of UID tags at the beginning and the end of the amplicon sequence [40]. We chose to introduce the 16 random nucleotides of the UID tag at the sequencing start site as the PGM semiconductor technology is significantly less accurate towards the end of the sequence [41]. We included a 4-nucleotide spacer as junction into the UID tag resulting in the N8-GATC-N8 ssUID layout of this study. Like this we address that the PGM indel rate increases in homopolymer stretches with their length [42], in particular when homopolymers are longer than 8nt [43]. While breaking potential homopolymer patterns within the UID, this design also reduces the number of mistakes during primer synthesis and allows to generate sets of primers with individual spacers that could be used to tag different experiments.

Nucleotide substitution errors are the most difficult to account for in HTS IG repertoire approaches and can critically falsify somatic hypermutation profiles [16,24]. They can originate from mixed events of adjacent insertions and deletions, which cannot be detected by the IMGT algorithm or are introduced as mistakes by the sequencing platform. UID barcoded RNA transcripts allow to address this problem [8,34,35,40]. B cells contain up to several thousands of identical IG RNA molecules that are each individually tagged by a UID [40,44]. Therefore, a HTS run provides a snapshot of the relative abundance of RNA transcripts [16]. Comparable to procedures used for identification of single nucleotide polymorphisms (SNP), single occurrences of nucleotide substitutions can be ruled out as artifacts and only transcripts above a certain copy threshold should be retained [44]. Our data show, that considering sequences with at least 2 copies in the final dataset improves the proportion of correct sequences by 0.7% to 99.5%. In this regard, as our sampling material are monoclonal hybridomas, all derived sequences (between 1,431 and 47,169) represent identical RNA molecules, making it stochastically more likely, that the same indel error appears several times. Thus, it is expectable, that the positive influence of excluding singletons would be even higher in bulk B cell derived datasets, where less sequences are derived from identical RNA molecule.

In conclusion, we have demonstrated that using our ssUID library preparation in combination with the IMGT database, the PGM sequencing platform can be efficiently used to assess murine IGH repertoires. Considering only consensus sequences with at least two copies improved the sequence quality considerably. Taken together, this approach allowed to obtain highly reliable IGH sequences, with more than 99% confidence in general and 99.9% confidence for the correct CDR3 sequences. The protocol and sample processing strategies described in this study will help to establish the benchtop-scale Ion Torrent sequencing technology of animal models in the field of immunoglobulin repertoire research.

## Materials and Methods

### RNA extraction

RNA was extracted with Trizol LS/chloroform (Thermo Fisher Scientific, Waltham, USA) method from seven monoclonal hybridoma cell lines (produced in house) with 10^6^ cells each. DNA was digested using the DNAfree kit (Thermo Fisher Scientific), RNA was further purified using Agencourt^®^ RNAclean XP beads (Analis, Suarlée, BE) and quantified on a NanoDrop^®^ Spectrophotometer (ND1000, Isogen Life Science, De Meern, NL). RNA was either directly used for library preparation or stored at -80°C.

### Reference sequences

Hybridoma cDNA transcripts were obtained using mouse constant region IgG primer (**suppl. table S2**) in a Superscript III (Thermo Fisher Scientific) reverse transcription following the manufacturer’s instructions for templates with high GC content. Transcripts were Sanger-sequenced (3100 Avant, Thermo Fisher Scientific) using constant region IgG and V-region primers (**suppl. table S2**). Forward and reverse sequences were aligned and submitted to IMGT V-QUEST (http://www.imgt.org, [45]) to verify the nucleotide sequence and to translate into amino acids. These sequences were subsequently used as reference sequences in alignments and artificial error insertion experiments.

### Datasets with artificial insertions and deletions

Artificial datasets were generated using the Biopieces indel_seq package (http://www.biopieces.org). For each of the original 7 hybridoma sequences, 2500 error-containing sequences were generated by combining 0-3 insertions and 0-3 deletions, obtaining a total of 37500 artificial sequences per hybridoma. For every set, indel-type and -position were determined by alignment to the original sequence to ensure homogenous error distributions. All artificial datasets were uploaded to IMGT HighV-QUEST and sorted by annotation: IMGT annotates correct sequences as productive. Sequences with a detected indel (frameshift, stop codon) are marked as “productive (see comment)” if the error can be corrected (referred to as “productive with detected errors”). Sequences with uncorrectable errors are classified as “unproductive”. If no fitting germline can be found sequences are marked as “unknown” or “no result” (referred to as “unknown/else”). The remaining indels on nucleotide level and amino acid changes were determined using the SeqAn library [46] in a custom-made C++ reference alignment program. For datasets with one insertion and one deletion (i1d1) the positions of the indels were determined by position-wise mismatch detection using a custom made Biopython [47] script. Upon detection, the nucleotide positions were returned and the process repeated with reverse complement sequences.

### Library preparation and HTS

Approximately 100ng (as determined by Nanodrop^®^) of total RNA per hybridoma was used for library preparation. We adapted the UID labeling method developed by Vollmers et al [40] to our PGM sequencing system (**suppl. Fig. S3**). RNA was reverse transcribed using Superscript III reverse transcriptase, according to the manufacturer’s instructions, using multiplex identifiers (MID) and UID tagged mouse constant region (IGHγ) primers elongated by partial PGM sequencing adapter pA (**suppl. Table S2**). The MID tag allowed multiplexing of several samples on one sequencing chip. The UID tag consists of two times 8 random nucleotides separated by a “GATC” spacer (N_8_-GATC-N_8_). With this UID tag each RNA molecule targeted by the primer is uniquely labeled (see [34,40] for detailed theoretical descriptions). The RT reaction mixtures were split into two equal second strand synthesis reactions using Phusion^®^ High-Fidelity DNA polymerase (NEB, Massachusetts, USA) with a mouse IGH V-region primer mix (**suppl. Table S2**). The reaction conditions were as follows: 98°C 2min, 50°C 2min, 72°C 10 min in a single cycle reaction. Both reaction aliquots were combined and purified twice using Agencourt^®^ AMPure^®^ XP beads (Analis) in a 1:1 (v/v) ratio to remove primer traces. Libraries were subsequently amplified with a Q5^®^ Hot Start High-Fidelity DNA polymerase (NEB) using the full-length Ion Torrent PGM sequencing adapters A and P1 as primers (**suppl. Table S2**) with the following conditions: 98°C for 1min, 20 cycles of 98°C for 10s, 65°C for 20s, 72°C for 30 seconds. Final elongation was done at 72°C for 2 min. Amplified libraries were purified twice using equal volumes of AMPure^®^ XP beads. Quality of the libraries as well as size of the amplicon and concentrations were determined using Agilent 2100 Bioanalyzer (Agilent Technologies, Diegem, BE) with the High Sensitivity DNA Kit (Agilent Technologies). 10 libraries were pooled equimolar on an Ion 316^TM^ Chip (Thermo Fisher Scientific) and sequenced on a PGM sequencer, with all quality trimming options disabled on the Torrent Suite^TM^ v4.0.2

### Data processing pipeline for the HTS datasets

Untrimmed raw reads were demultiplexed by their MIDs, retaining only sequences containing the full UID primer sequence for further analysis, with no mismatches allowed. The UID sequence was extracted and categorized in relation to the starting position of the detected primer including the GATC spacer and stored in the sequence identifier. After clipping the MID, UID and constant region primer, the trimmed reads were quality controlled (80% of the bases Phred-like quality score above 20) and grouped into UID families. Using pagan-msa [48], a consensus sequence was generated for each UID-family containing more than 2 members. Afterwards, reverse primers were identified with up to 2 mismatches and clipped. Subsequently, sequences were collapsed to unique reads, storing counts in the read identifier, and uploaded to IMGT for error detection, correction, annotation and translation into amino acids. Post-IMGT datasets were separated into four categories (“productive”, “productive with detected errors”, “unproductive” and “unknown/else”) and processed separately. Data processing was performed using custom-made Python scripts (Python v2.7) employed in a parallelizing bash wrapper script using gnu-parallel [49] and the Biopieces framework (http://www.biopieces.org/).

### Graphs and statistics

All graphs and statistical analyses were performed using R base packages or GraphPad Prism 6. Average numbers are reported as mean ± standard deviation (SD) unless specified otherwise.

## Figure legends

**Figure S1: Indel positions for mixed i1d1 datasets of hybridomas 2-7.** The distances between the first and second indel before correction in the i1d1 dataset of hybridomas 2-7 are shown as scatterplots. Dotted lines indicate positions of IMGT junctions. Productive sequences with detected indels are shown in grey, unproductive sequences are shown in dark grey. Sequences without detected errors are shown in black.

**Figure S2: Additional artificial indel set alignments.** Indel positions are shown before and after IMGT error correction for artificially falsified Hybridoma 1 sequences separated by productivity. The indels for the datasets i1d2, i1d3, i2d1, i2d2, i2d3, i3d1, i3d2, i3d3, i0d2, i0d2 are shown per nucleotide position as line plot (smoothened over 4 neighbors). The grey area marks the IGH VDJ junction.

**Figure S3. 3-step PGM ssUID sequencing library preparation.** (A) In a first step, purified mRNA is used in a Superscript III reverse transcription. The Primer for the reverse transcription is specific for the murine IG C region and elongated by an MID for sample multiplexing as well as a UID consisting of 2x 8 random nucleotides (N8) separated by a 4-nucleotide spacer (‘GATC’). The primer ends with the partial PGM sequencing adapter pA. (B) In the second step, a mix of 26 IG VH region targeting primers (elongated by the partial PGM sequencing adapter pP1) is used in a single cycle PCR reaction to avoid amplification. The product of this reaction is purified twice with Agencourt^®^ AMPureXP beads to remove the VH primers from the reaction mixture. (C) In the final step, the purified reaction mixture is amplified using the full-length P1 and A adapters as primers in a 20 cycle PCR reaction. The product is as well purified twice to obtain the ssUID-tagged sequencing library.

## Abbreviations

CDR3: complementary determining region 3
HTS: high-throughput sequencing
IG: immunoglobulin
IGH: immunoglobulin heavy chain
IMGT: ImMunoGeneTics
indel: insertions and deletions of nucleotides
MID: multiplex identifier
nt: nucleotide
PGM: (Ion Torrent) Personal Genome Machine
UID: Unique (molecular) identifier
ssUID: single side unique molecular identifier

## Authors Contribution

J-P.B. designed research, cultivated hybridomas, performed library preparation, developed bioinformatics approaches, performed data processing, interpreted data and wrote the manuscript. W.J.F. and O.H. supported and developed bioinformatics approaches and performed data processing. A.R.S.X.D designed research and interpreted data. A.W-B. developed and wrote the raw data processing bioinformatics pipeline. R.S. performed Ion Torrent PGM sequencing. A.B. designed research, supervised work, assisted library preparation and hybridoma cultivation and interpreted data C.P.M. supervised work, provided important intellectual input and interpreted data. All authors have read and corrected the manuscript.

## Acknowledgements

J.-P.B. was supported by the Aides à la Formation-Recherche (AFR) individual PhD grant of the Fonds National de la Recherche of Luxemburg (www.fnr.lu, grant 7039209). We would like to thank Josiane Kirpach for independently verifying the library preparation protocol and Fleur A.D. Leenen for critically revising the manuscript.

